# Ancestral acetylcholine receptor β-subunit forms homopentamers that prime before opening spontaneously

**DOI:** 10.1101/2022.01.12.475990

**Authors:** Christian J. G. Tessier, R. Michel Sturgeon, Johnathon R. Emlaw, Gregory D. McCluskey, F. Javier Pérez-Areales, Corrie J. B. daCosta

## Abstract

Human adult muscle-type acetylcholine receptors are heteropentameric ion channels formed from two α-subunits, and one each of the β-, δ-, and ε-subunits. To form functional channels, the subunits must assemble with one another in a precise stoichiometry and arrangement. Despite being different, the four subunits share a common ancestor that is presumed to have formed homopentamers. The extent to which the properties of the modern-day receptor result from its subunit complexity is unknown. Here we show that a reconstructed ancestral muscle-type β-subunit can form homopentameric ion channels. These homopentamers open spontaneously and display single-channel hallmarks of muscle-type acetylcholine receptor activity. Our findings demonstrate that signature features of muscle-type acetylcholine receptor function are independent of agonist, and do not necessitate the complex heteropentameric architecture of the modern-day receptor.

## Introduction

Ligand-gated ion channels convert chemical signals into electrical impulses by coupling the binding of small molecules to the opening of an ion-conducting transmembrane pore (1). Of the many types of ligand-gated channels, the superfamily of pentameric ligand-gated ion channels (pLGICs) is the largest and most structurally and functionally diverse (2). Formed from five identical or homologous subunits arranged around a central ion-conducting pore, pLGICs are found in almost all forms of life (3). In prokaryotes, pLGICs appear to be exclusively homopentameric, while in eukaryotes both homo- and heteropentameric channels are common. Humans express more than forty different pLGIC subunits, which are subdivided based on whether they form cation-selective channels activated by acetylcholine or serotonin (5-hydroxytryptamine; 5-HT_3_), or anion-selective channels activated by GABA (γ-aminobutyric acid) or glycine. This repertoire of pLGIC subunits, combined with their ability to form homo- and heteropentamers, leads to a wealth of structural and functional diversity, presumably to meet a variety of synaptic needs (4).

The archetypal pLGIC is the heteropentameric muscle-type nicotinic acetylcholine receptor (AChR), with the human adult form composed of two α-subunits, and one each of the β-, δ-, and ε-subunits (Fig. 1)(5, 6). To gain insight into the structure, function, and evolution of the AChR, we have employed an ancestral reconstruction approach (7–10). Previously we resurrected a putative ancestral AChR β-subunit. Referred to as “β_Anc_”, this subunit differed from its human counterpart by 132 amino acids (i.e. approximately 30% of the total amino acid sequence), yet was able to substitute for the human β-subunit, and also supplant the human δ-subunit, forming functional hybrid ancestral/human AChRs (7). These hybrid AChRs were ternary mixtures, containing two β_Anc_ subunits, two human α-subunits, and one human ε-subunit. A concatameric construct confirmed that the two β_Anc_ subunits resided next to each other, occupying positions in the heteropentameric complex usually reserved for the human β- and δ-subunits (Fig. 1) (7). Regardless of whether they bind agonist or not, all pLGIC subunits have both principal (+) and complementary (−) subunit interfaces. To sit next to each other, the (+) and (−) interfaces of any two neighbouring subunits must be structurally compatible. Thus, for β_Anc_ to replace both the human β- and δ-subunits, the (+) and (−) interfaces of β_Anc_ must be compatible with each other, raising the possibility that multiple β_Anc_ subunits could coassemble to form β_Anc_ homomers (Fig. 1).

**Figure 1:**
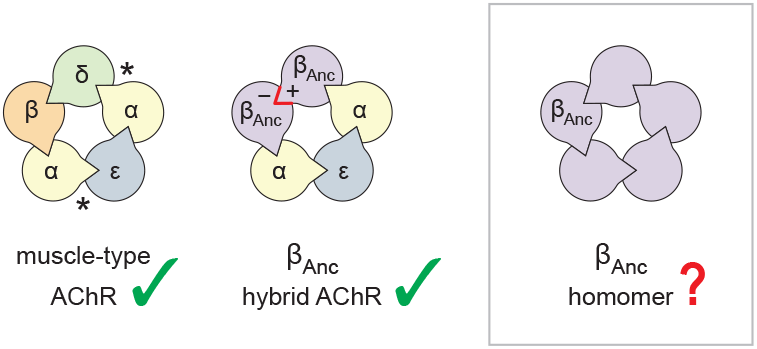
Subunit composition of heterologously expressed acetylcholine receptors. Subunit stoichiometry and arrangement of the human adult muscle-type acetylcholine receptor (left), where the agonist-binding sites at the α-δ and α-ε subunit interfaces are indicated with asterisks (*). A reconstructed ancestral β-subunit (β_Anc_; purple) forms hybrid acetylcholine receptors (middle) where β_Anc_ substitutes for the human β-subunit (β; orange) and supplants the human δ-subunit (δ; green). The principal (+) and complementary (−) interfaces of β_Anc_ must be compatible for two β_Anc_ subunits to sit side-by-side (red highlight), which predicts that homomers formed from multiple β_Anc_ subunits should be possible (right, boxed).

Here, using single-channel measurements, we demonstrate that β_Anc_ readily forms homopentameric channels, which open spontaneously in the absence of acetylcholine. This spontaneous activity displays hallmarks of the muscle-type AChR, including steady-state single-channel burst behaviour. These findings demonstrate how fundamental characteristics of AChR activation are independent of agonist, and not a result of the complex heteropentameric architecture of the muscle-type AChR. Finally, an alternate ancestral β-subunit, reconstructed using a phylogenetic tree that matched the accepted species relationships (7), and which only shared ~85% identity with β_Anc_, revealed that these unexpected characteristics of β_Anc_ are robust to phylogenetic uncertainty, and thus deeply embedded within AChR β-subunit structure and evolutionary history.

### Experimental Procedures

#### Materials

2-[(2,6-Dimethylphenyl)amino]-*N*,*N*,*N*-trimethyl-2-oxoethaniminium chloride (QX-222) was purchased from Tocris Bioscience. All other chemicals, including acetylcholine chloride, were purchased from Sigma-Aldrich.

#### Molecular Biology

cDNAs of human muscle-type AChR subunits (⍺1, β1, δ, and ε) in the pRBG4 plasmid were provided by Steven M. Sine (Mayo Clinic), while cDNAs encoding β_Anc_ and an alternate ancestral β-subunit (β_AncS_) were reconstructed and cloned into pRBG4 as described previously (7, 10). Mutations to produce the low-conductance variant of β_Anc_ (β_AncLC_ = E420R, D424R, E428R) were introduced by inverse PCR (11). Sanger sequencing confirmed the entire reading frame for all constructs.

#### Mammalian Cell Expression

Combinations of human and ancestral subunit cDNAs were transfected into BOSC 23 cells (12). Cells were maintained in Dulbecco’s modified Eagle’s medium (DMEM; Corning) containing 10% (vol/vol) fetal bovine serum (Gibco) at 37°C, until they reached 50-70% confluency. Cells were then transfected using calcium phosphate precipitation, and transfections terminated after three to four hours by exchanging the medium. All experiments were performed one day post transfection (between 16-24 hours after exchanging the medium). A separate plasmid encoding green fluorescent protein was included in all transfections to facilitate identification of transfected cells.

#### Single-Channel Patch Clamp Recordings

Single-channel patch clamp recordings were performed as previously described (13). Recordings from BOSC 23 cells transiently transfected with cDNAs encoding wild-type, ancestral, or low-conductance subunits, were obtained in a cell-attached patch configuration. All recordings were obtained with a membrane potential of –120 mV, with room temperature maintained between 20-22 °C. The external bath solution contained 142 mM KCl, 5.4 mM NaCl, 0.2 mM CaCl_2_ and 10 mM 4-(2-hydroxyethyl)-1-piperazineethanesulfonic acid (HEPES), adjusted to pH 7.40 with KOH. The pipette solution contained 80 mM KF, 20 mM KCl, 40 mM K-aspartate, 2 mM MgCl_2_, 1 mM ethylene glycol-bis(β-aminoethyl ether)-N,N,N′,N′-tetraacetic acid (EGTA), and 10 mM HEPES, adjusted to a pH of 7.40 with KOH. Acetylcholine and QX-222 were added to pipette solutions at their desired final concentrations and stored at –80 °C. Patch pipettes were fabricated from type 7052 or 8250 nonfilamented glass (King Precision Glass) with inner and outer diameters of 1.15 and 1.65 mm, respectively, and coated with SYLGARD 184 (Dow Corning). Prior to recording, electrodes were heat polished to yield a resistance of 5-8 MΩ. Single-channel currents were recorded on an Axopatch 200B patch clamp amplifier (Molecular Devices), with a gain of 100 mV/pA and an internal Bessel filter at 100 kHz. Data were sampled at 1.0 μs intervals using a BNC-2090 A/D converter with a National Instruments PCI 6111e acquisition card and recorded by the program Acquire (Bruxton).

#### Dwell Time and Kinetic Analysis

Single-channel detections were performed using the program TAC 4.3.3 (Bruxton). Data were analyzed with an applied 10 kHz digital Gaussian filter. Opening and closing transitions were detected using the 50% threshold crossing criterion, and open and closed dwell duration histograms were generated within the program TACfit 4.3.3 (Bruxton). Histograms were visually fit with a minimum sum of exponential components. From the closed duration histograms, the intersection of the slowest activation and fastest inactivation/desensitization components was taken as the critical closed duration (τ_crit_) (14, 15). Closings longer than τ_crit_ (corresponding to inter-burst closings) were removed from analysis. Events were imported into R using *scbursts* (16), where individual durations were corrected for instrument risetime (17), and bursts were defined by a τ_crit_ (15). Bursts with fewer than three events were omitted from further analysis. The probability of being open within a burst (i.e. burst *P*_open_) was calculated for each burst, and bursts with a *P*_open_ that did not fit within the normal distribution were removed using *extremevalues* (18). The distribution of burst *P*_open_ was then fit with a Gaussian distribution, and bursts within two standard deviations from the mean were used for further kinetic analysis (16). A user-defined kinetic scheme (see Figure S2 for β_Anc_, and the modified del Castillo and Katz scheme (19) for the human adult acetylcholine receptor in Figure S3) was fit to the sequence of single-channel dwells in the global dataset using maximum likelihood implemented within MIL (QUB suite, State University of New York, Buffalo, NY). With a user-defined dead time of 18.83 μs, MIL corrected for missed events, estimated model parameters by maximum likelihood, and gave standard errors of the estimated parameters (See Supplemental Tables S1, S2, and S3) (20).

#### Electrical Fingerprinting

High-(HC) and low-conductance (LC) variants of β_Anc_ were transfected at 1:4 and 4:1 (HC:LC) cDNA ratios. Transfections and single-channel recordings were performed as described above. For detections, data were filtered to 1 kHz, and bursts defined by a uniform τ_crit_ of 2 ms imposed upon all recordings. Using the program TAC 4.3.3. (Bruxton), amplitudes of single-channel bursts were measured as the difference between open- and closed-channel currents. Amplitudes of individual bursts were pooled from separate recordings to generate event-based amplitude histograms (EBAHs; Fig. 5), which were fit with Gaussian distributions within TACfit (Bruxton).

## Results

Our first inkling that a reconstructed ancestral β-subunit (β_Anc_) may be able to form homomeric channels came from heterogeneity in single-channel recordings acquired after alterations to our original transfection protocol. Typical cotransfection of cDNAs encoding human α-, δ-, and ε-subunits with a cDNA encoding β_Anc_ leads to robust cell surface expression of β_Anc_-containing hybrid AChRs (10). In an attempt to lower overall AChR expression with the purpose of facilitating single-channel analysis, we reduced the total amount of subunit cDNA in our transfections (~6-fold), while maintaining the same 2:1:1:1 subunit cDNA ratio (by weight; α:β_Anc_:δ:ε). Reducing the amount of cDNA lowered overall AChR expression as expected, but also led to a previously unseen heterogeneity in our patches (Fig. 2A). Instead of a single population of channels with a uniform amplitude of ~10 pA, and a burst behavior indicative of β_Anc_-containing AChRs (Fig. 2A; left inset), we also observed a second class of channels with a different kinetic signature, and an increased amplitude of ~16 pA (Fig. 2A; right inset). A similar trend was not observed when cDNA encoding the wild-type human β-subunit was cotransfected instead of β_Anc_ (Fig. S1), indicating that the ancestral β-subunit was the source of the heterogeneity. Consistent with this, lowering the proportion of β_Anc_ cDNA in the transfection mixture reduced the fraction of high amplitude channels (Fig. 2B,D), while transfecting exclusively with β_Anc_ cDNA resulted in patches where all channel openings had a uniformly high amplitude (Fig. 2C,D). This demonstrated that when transfected alone, β_Anc_ forms functional ion channels.

**Figure 2:**
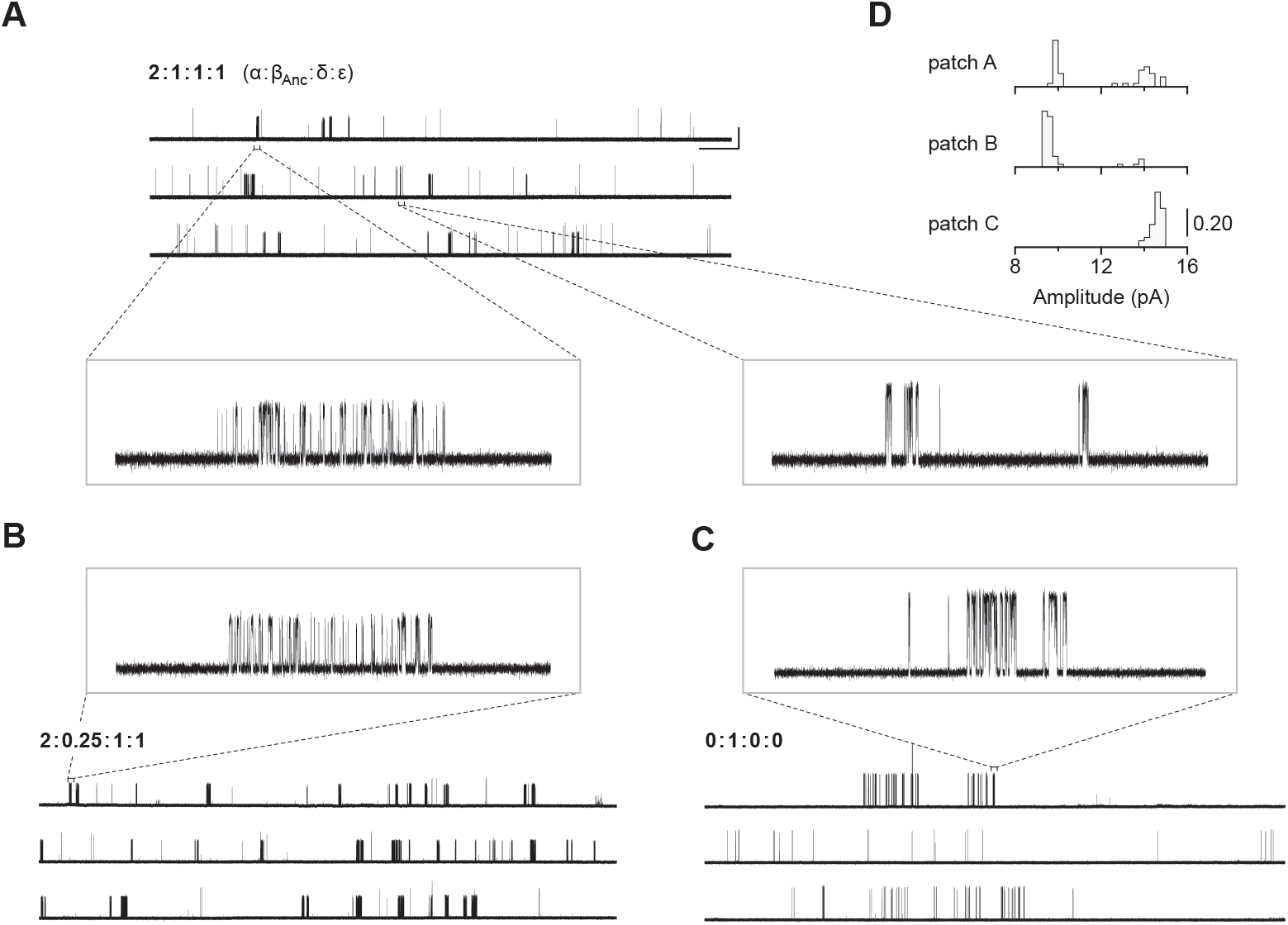
Single-channel recordings of β_Anc_-containing channels. (A) Representative continuous recording from a cell-attached patch where cells were transfected with cDNAs encoding human muscle-type α-, δ-, and ε-subunits, and an additional cDNA encoding β_Anc_ at a cDNA ratio of 2:1:1:1 (α:β_Anc_:δ:ε). (B) Same as in panel “A”, but where cells were transfected with an altered 2:0.25:1:1 cDNA ratio, making the β_Anc_ subunit limiting, or (C) where cells were transfected with only the cDNA encoding β_Anc_. In all cases openings are upward deflections, in the presence of 30 μM acetylcholine, and with an applied voltage of –120 mV. Continuous recordings are digitally filtered to 5 kHz, and the scale bar (2 s, 10 pA) in “A” applies to “B” and “C”. Insets are digitally filtered to 10 kHz, with boxes representing scale bars (300 ms, 25 pA). (D). Event-based amplitude histograms for single-channel bursts from each of the patches shown in “A”, “B” and “C”. In each case, the height of the bins was normalized to the total number of bursts in each patch (A: 40; B: 50; C: 41), with the scale bar representing the indicated fraction (0.20) of the total bursts.

The traces in Figure 2 were recorded in the presence of agonist (30 μM acetylcholine). In the human adult muscle-type AChR, the agonist-binding sites are located at the α-δ and α-ε interfaces, and the β-subunit is the only subunit that does not participate directly in agonist binding (Fig. 1) (21). We were therefore surprised to see single-channel activity in patches from cells transfected exclusively with β_Anc_, as channels formed from a muscle-type β-subunit alone would not be expected to have intact agonist-binding sites. To determine if the activity of β_Anc_-alone channels was dependent upon acetylcholine, we recorded single-channel activity in the absence of acetylcholine (Fig. 3). When no acetylcholine was present, patches from cells that were transfected exclusively with β_Anc_ cDNA still displayed single-channel activity, indicating that β_Anc_-alone channels open spontaneously under these conditions (Fig. 3A). Furthermore, spontaneous activity occurred as bursts of closely spaced openings, separated by brief closings (Fig. 3B)(14). The briefest of these intervening closings were reminiscent of classic “*nachschlag* shuttings” (Fig. 3B, “*ii*” in inset), observed in early patch clamp recordings from frog end-plate nicotinic receptors, and originally thought to relate to agonist efficacy (22). Thus, despite being homomeric and lacking agonist-binding sites, β_Anc_-alone channels display single-channel hallmarks of the muscle-type AChR.

**Figure 3:**
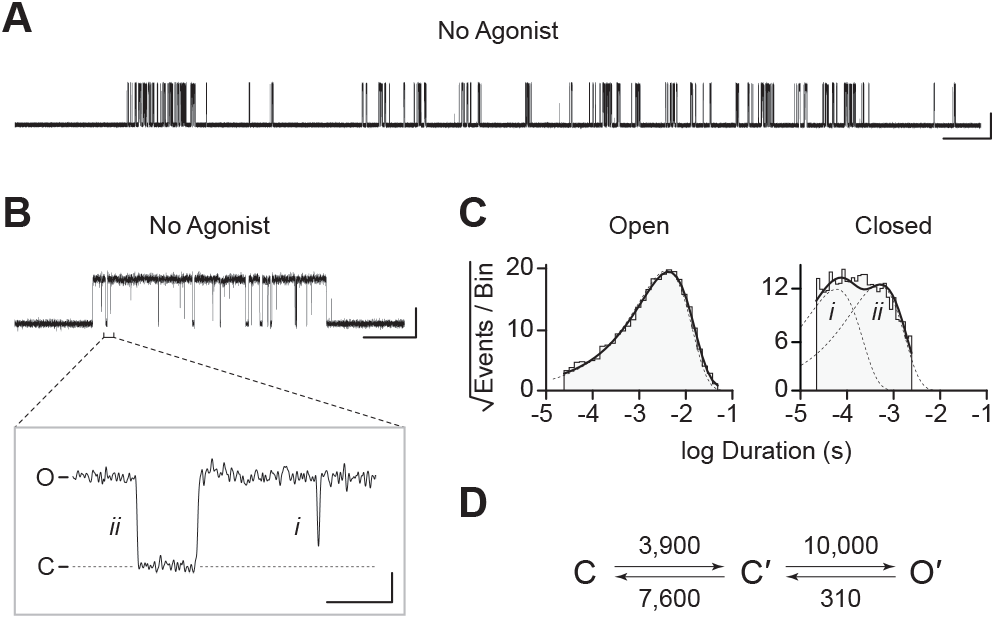
Spontaneous single-channel openings of β_Anc_ homomers. (A) Representative continuous recording of a cell-attached patch from cells transfected with a single cDNA encoding β_Anc_. Recording was made in the absence of acetylcholine and at an applied voltage of –120 mV. Data was digitally filtered to 5 kHz (scale bar = 2 s, 10 pA). (B) Single burst of openings from a homomeric β_Anc_ channel, shown digitally filtered to 10 kHz (scale bar = 25 ms, 10 pA). Inset depicts (*i*) brief and (*ii*) long closings within bursts, where the former (*i*) are reminiscent of “*nachschlag* shuttings” (scale bar = 1 ms, 5 pA). (C) Open and closed dwell duration histograms for the representative patch depicted in “B”. Individual exponential components determined manually (dashed lines) and kinetic fits from MIL (solid lines) are overlaid. Global kinetic fitting was performed on three individual recordings, from two separate transfections. (D) The single-channel data fit a three-state scheme (Scheme 1), where C, C′, and O′ correspond to closed, closed-primed, and open-primed states. Rate constants are shown above and below corresponding arrow, with error estimates provided in Table S1.

To gain insight in the spontaneous activity of β_Anc_-alone channels, we performed kinetic analysis of our single-channel data. First, we determined a critical closed duration (τ_crit_) to define bursts arising from a single ion channel. Then we determined the minimum number of components in our apparent open and closed dwell duration histograms by fitting each with a sum of exponentials. Open duration histograms were fit well by a single exponential component, while closed duration histograms required at least two components (Fig. 3C). This suggested that a minimal scheme with a single open state and two closed states is necessary and sufficient to describe the spontaneous activity of β_Anc_-alone channels. Based on this, we then fit the sequence of single-channel dwells using the three possible kinetic schemes, two linear and one cyclic, comprising a single open state and two closed states (Fig. 3D; Fig. S2). As a control we also fit a simplified two-state scheme, where a single open state was connected to a single closed state, which, based on the relatively poor fit of the closed durations, confirmed that inclusion of a second closed state was justified (Fig. S2). Overlaying the resulting fits on top of duration histograms revealed that each of the possible three-state schemes fits the observed dwells equally well, thus discriminating between the possible kinetic schemes is not trivial (Fig. S2). We settled upon the simple linear scheme, where β_Anc_-alone channels transition from a closed state, “C”, to an intermediate closed state, “C′”, before opening to “O′”. The form of this scheme, with an intermediate closed state that precedes channel opening, is guided by models of AChR activation that include a single “flipping” or multiple “priming” steps (13, 23, 24). Given that for β_Anc_-alone channels there is a single intermediate closed state that precedes channel opening, we refer to this state in our scheme as “flipped” or “primed”.

In the presence of acetylcholine, the single-channel current traces appeared different (Fig. 4A). As the concentration of acetylcholine increased from 10 to 100 μM, there was a progressive decrease in the duration of openings, as well as a reciprocal increase in the number of short-lived closings within each burst (Fig. 4A). This can be observed as a leftward shift in the open duration histograms as the concentration of acetylcholine is increased. This single-channel behavior is a hallmark of open-channel block, a ubiquitous property of AChR agonists, including acetylcholine (25). Consistent with this blockage profile, the same trend, albeit with longer-lived blocking events, was observed with the well-characterized AChR open-channel blocker, QX-222 (Fig. 4B)(26–28).

**Figure 4:**
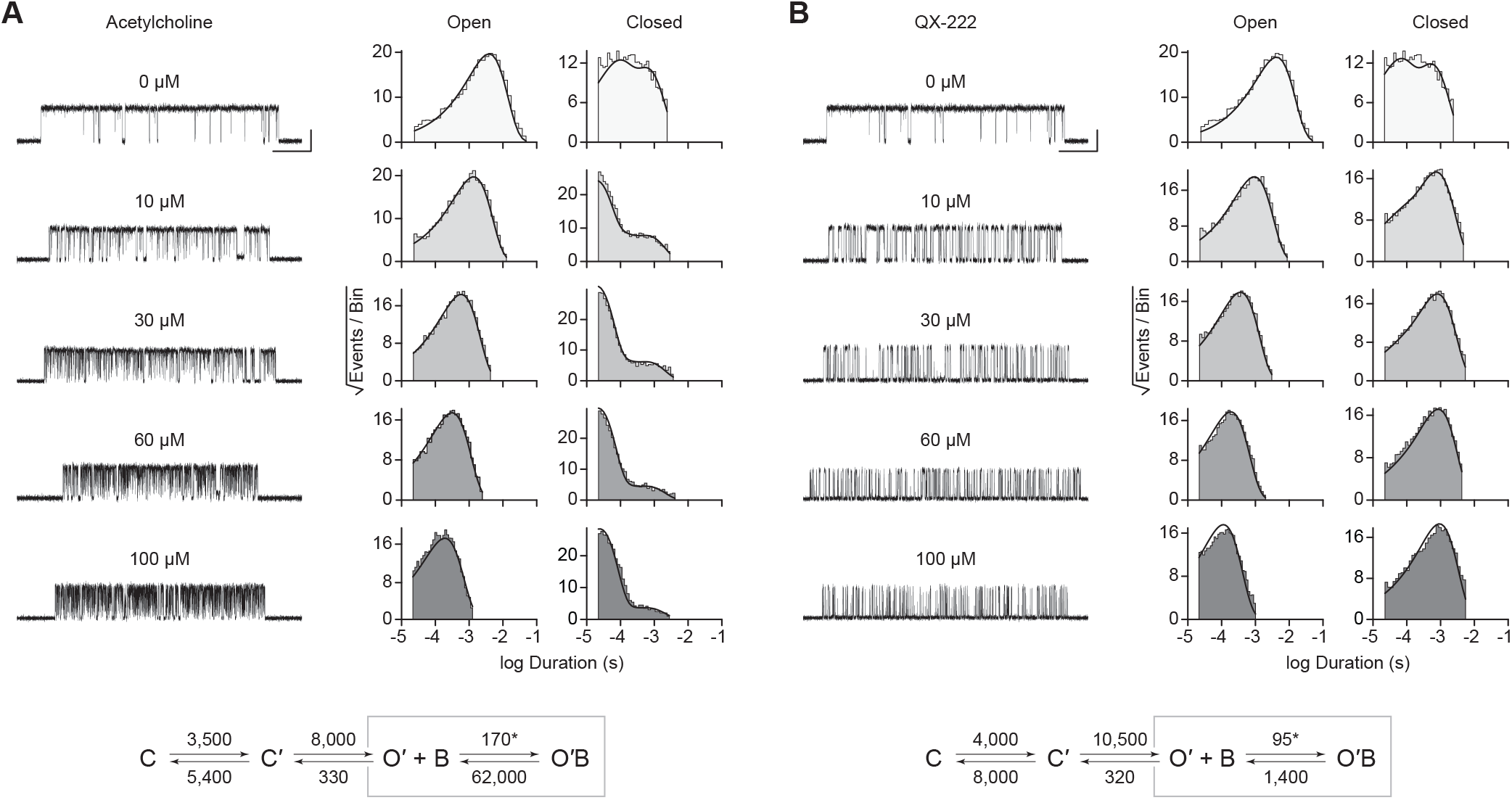
Open-channel block of β_Anc_ homomers by acetylcholine and QX-222. Representative single-channel activity of β_Anc_ homomers in the presence of increasing concentrations of (A) acetylcholine, and (B) QX-222. Openings are upward deflections. Recordings were obtained with an applied voltage of –120 mV. Data were filtered to 10 kHz (scale bars = 25 ms, 10 pA; applies to “A” and “B”). The sequence of dwells from each data set, encompassing the full concentration range of the blocker, was globally fit to the same three-state scheme used for β_Anc_, where an additional fourth state corresponding to the open/blocked channel was added (Scheme 2). Global kinetic fits were performed on three individual recordings for each concentration of blocker, from at least two separate transfections, corresponding to 15 total patches for each global fit. Note that the recordings in the absence of blocker are the same for each data set. Rate constants with error estimates are presented in Table S1.

The kinetics of open-channel block are determined by interactions between the blocking molecule and residues that line the channel pore in the open state, and therefore provide indirect structural insight into the open state. To compare the open state structures of β_Anc_-alone homomers with wild-type AChRs, we determined the kinetics of acetylcholine and QX-222 block for both types of channels (Fig. 4; Fig. S3; Fig. S4). To fit our β_Anc_-alone single-channel data recorded in the presence of a blocker, we introduced an additional open, but blocked (i.e. non-conducting) state connected to our open state, where the forward rate of block was dependent upon the concentration of the blocking molecule (Fig. 4A,B). We then globally fit each of our β_Anc_-alone data sets encompassing between 0 and 100 μM acetylcholine or QX-222. Initially, we restricted the rates of the core (C-C′-O′) scheme to those inferred in the absence of blocker, however allowing all parameters to be estimated, led to negligible changes in the inferred rates of block. For β_Anc_-alone and wild-type channels, the rates of acetylcholine and QX-222 block were comparable (see Tables S1,S2), suggesting that the structure of the open pore in the two types of channels is similar.

Given that β_Anc_-alone channels are expressed in the absence of other AChR subunits, a reasonable hypothesis is that they are homopentamers. To determine the subunit stoichiometry of β_Anc_-alone channels, we employed a single-channel electrical fingerprinting strategy, where mutations altering unitary conductance are used to count the number of individual β_Anc_ subunits in β_Anc_-alone channels. A similar strategy has been employed with tetrameric potassium channels (29), and other pLGICs (30), including both the homopentameric α7 AChR (31–33) and the heteropentameric muscle-type AChR (7). The approach relies on identifying high-conductance (HC) and low-conductance (LC) variants of the β_Anc_ subunit, and then co-expressing them to reveal a number of amplitude classes. Openings in each amplitude class originate from channels incorporating the same ratio of HC to LC subunits, and based on the total number of amplitude classes, the number of β_Anc_ subunits within β_Anc_-alone channels can be inferred.

When β_Anc_ is expressed alone, the resulting channels exhibit a single, uniform amplitude, distributed around a mean of ~16 pA (Fig. 5A,D), making the wild-type β_Anc_ subunit an ideal high-conductance (HC) subunit for electrical fingerprinting. To identify a low-conductance (LC) variant of β_Anc_, we took advantage of a structural feature inherent to eukaryotic pLGICs: as conducting ions exit the channel’s transmembrane pore, they are obliged to pass through one of five portals in the cytoplasmic domain (21, 34). Framed by charged or polar residues from each subunit, these portals influence single-channel conductance. Mimicking the homologous 5-HT_3A_ receptor, which has an unusually low single-channel conductance (35–37), we substituted three arginine residues (E420R, D424R, E428R) into this region of β_Anc_. When β_Anc_ harbouring three arginines in this region was transfected by itself, the resulting channels exhibited a reduced single-channel amplitude centered around ~1-2 pA (Fig. 5B,D). With its markedly reduced amplitude, β_Anc_ harbouring three arginine residues is a suitable LC subunit for electrical fingerprinting.

**Figure 5:**
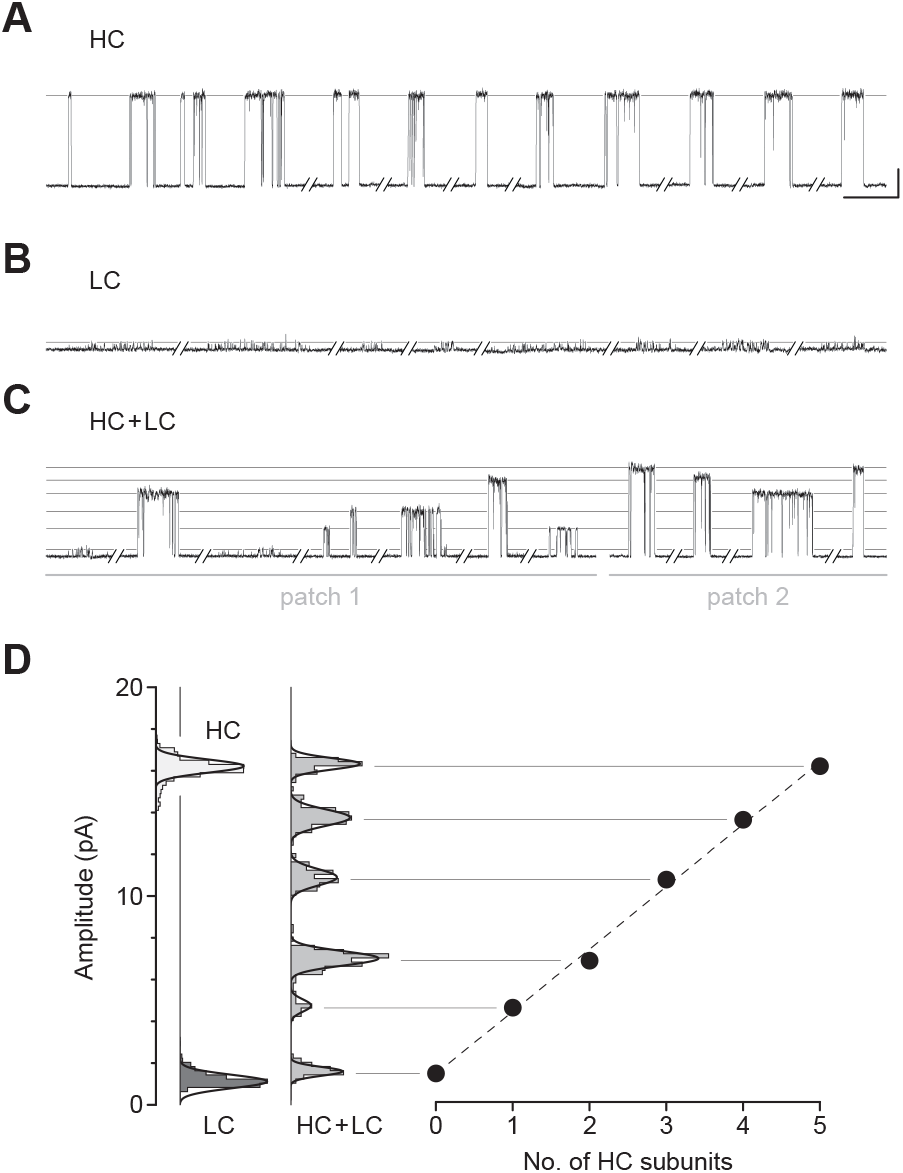
Electrical fingerprinting to determine subunit stoichiometry of β_Anc_ homomers. Representative single-channel activity from cells transfected with (A) cDNA encoding the wild-type high-conductance (HC) β_Anc_ subunit, or (B) a mutant low-conductance (LC) β_Anc_ variant harbouring substitutions that reduce single-channel amplitude. (C) Cotransfection of cDNAs encoding HC and LC β_Anc_ variants led to patches (two shown) with heterogeneous amplitudes. (D) The amplitudes segregate into six well-defined amplitude classes (combined from the two patches in “C”), where the highest and lowest amplitude classes match that of the all-HC and all-LC classes, respectively. Plot of the mean amplitude of each class as a function of the presumed number of incorporated HC subunits (error bars = standard deviations of the mean but are smaller than the points themselves). Recordings were obtained with an applied voltage of –120 mV, and traces were digitally filtered to 1 kHz to facilitate amplitude detection (scale bar = 25 ms, 10 pA; applies to “A”, “B”, and “C”).

When cDNAs encoding HC and LC variants of β_Anc_ were transfected together, a variety of single-channel amplitudes were observed in each patch (Fig. 5C). The relative proportion of channels with high versus low amplitude could, to some degree, be tuned by the ratio of HC to LC β_Anc_ cDNA used for transfection (Fig. S5). Constructing event-based amplitude histograms, and pooling amplitudes from more than one recording, revealed that the amplitudes segregated into as many as six amplitude classes, with the highest and lowest amplitude classes matching that of the HC and LC forms of β_Anc_-alone channels. The difference in amplitude between successive classes was somewhat regular, demonstrating five approximately equal contributions to single-channel conductance (Fig. 5D), consistent with the hypothesis that β_Anc_-alone channels are homopentamers.

As noted previously, reconstruction of β_Anc_ was based upon a molecular phylogeny that diverged from the accepted species phylogeny (7–10). Reconstruction of ancestral protein sequences is based upon a best-fit model of amino acid evolution, a multiple sequence alignment, as well as a phylogenetic tree relating the sequences within the alignment (38). We therefore wondered if the ability of β_Anc_ to form spontaneously opening homomers was an artefact of the discordant tree used to reconstruct it. To test this, we took advantage of an alternate ancestral β-subunit, called “β_AncS_”, whose reconstruction was based upon a molecular phylogeny that matched the accepted species phylogeny (7). Despite 67 substitutions and 6 indels relative to β_Anc_, when expressed alone, β_AncS_ still formed homomeric channels that opened in bursts in the absence of acetylcholine (Fig. 6). This demonstrated that the ability of β_Anc_ to form homomers that spontaneously open is not an artefact of the phylogeny used to reconstruct it. Instead, this surprising ability of β_Anc_ is robust to the phylogenetic uncertainties inherent in ancestral sequence reconstruction, as well as substantial variation in the amino acid sequence of the reconstructed β-subunits.

**Figure 6:**
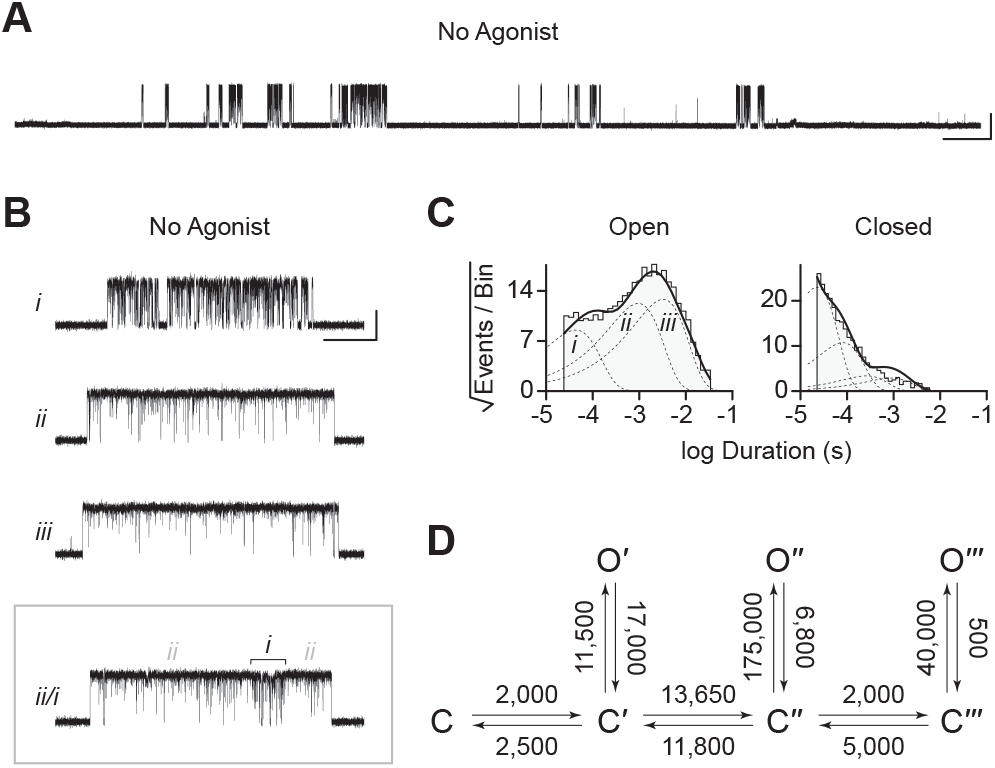
Spontaneous single-channel openings of homomers formed from an alternate ancestral β-subunit (β_AncS_). (A) Representative continuous recording of a cell-attached patch from cells transfected with a single cDNA encoding β_AncS_. Recording was made in the absence of acetylcholine, at an applied voltage of –120 mV, where spontaneous openings are upward deflections. Data was digitally filtered to 5 kHz (scale bar = 2 s, 10 pA). (B) Bursts from homomeric β_AncS_ channels, each exhibiting one of three different types (*i*, *ii*, *iii*) of openings (scale bar = 25 ms, 10 pA). The boxed burst at the bottom is an example of a single burst that contains more than one type of opening (*ii/i*). (C) Open and closed dwell duration histograms for the representative patch depicted in “B”. Individual exponential components determined manually (dashed lines) and kinetic fits from MIL (solid lines) are overlaid. Global kinetic fitting was performed on three individual recordings, from two separate transfections. The exponential components (*i*, *ii*, *iii*) in the open duration histogram correspond to the different types of openings observed within the bursts in panel “B”. (D) The three-state scheme in Figure 3 (Scheme 1) was expanded to include additional priming steps (“singly”, “doubly”, and “triply” primed), each with their own connected open state (Scheme 3). Rate constants are shown, with error estimates provided in Table S3.

While β_AncS_ forms homomers that open spontaneously, inspection of the β_AncS_ single-channel activity revealed additional complexity not seen with the original β_Anc_ subunit. For β_AncS_, single-channel bursts were heterogeneous, displaying at least three distinct kinetic behaviours (Fig. 6B). In some cases, the kinetic behaviour changed within a burst, demonstrating that the different kinetics were possible within the same channel (Fig. 6B; boxed). This heterogeneity was also reflected in apparent open and closed duration histograms, with each displaying a minimum of three exponential components (Fig. 6C). The increased number of exponential components indicated additional open and closed states relative to β_Anc_, and thus that the three-state scheme used to fit β_Anc_ was insufficient to describe the spontaneous activity of β_AncS_. To account for the additional states, we expanded our original scheme to include two additional “priming” steps, where openings could occur from one of three primed states (Fig. 6D). This scheme, with multiple priming steps, builds directly upon the one used to fit muscle-type AChRs that had been engineered to open spontaneously (24). Given the heterogeneity of the β_AncS_ single-channel activity, and the complexity of this scheme, we caution against over interpretation of the inferred rates. Nevertheless, we note that in accord with the muscle-type AChR, the equilibrium gating constants appear to increase (Table S3; compare Θ_1_, Θ_2_, Θ_3_), and thus the open states become more and more favoured, for each successive priming step. In any case, the fits suggest that a scheme of this form, with multiple stages of priming, is adequate to describe the complex spontaneous single-channel activity of β_AncS_ homomers.

## Discussion

We have shown that a reconstructed ancestral acetylcholine receptor β-subunit (β_Anc_) readily forms homopentameric channels. At first glance, this may seem unexpected. Modern-day muscle-type β-subunits do not appear to form homopentamers, and instead appear fully entrenched within heteropentameric muscle-type AChRs. Since reconstruction of β_Anc_ was informed almost exclusively by modern muscle-type β-subunits, there was no reason to expect that β_Anc_ would behave differently (10). However, as mentioned previously, β_Anc_ can replace both the human muscle-type β- and δ-subunits in hybrid ancestral/human AChRs (7). For this to be possible the principal (+) and complementary (−) subunit interfaces of β_Anc_ must be compatible with each other (Fig. 1), leading to the logical hypothesis, confirmed here, that β_Anc_ can form homopentamers. This ability to revert the muscle-type β-subunit, which is entrenched in a heteropentamer, back to a subunit capable of forming homopentamers, attests to the homopentameric origins of all pentameric ligand-gated ion channel subunits (39).

We have also shown that β_Anc_ homopentamers open spontaneously. This is surprising. What is even more surprising is that the spontaneous single-channel activity of β_Anc_ homopentamers resembles that of the agonist-activated muscle-type AChR. Some of the first single-channel recordings of frog AChRs activated by agonists revealed that agonist-activated openings occur in quick succession, as bursts of activity originating from the same channel (40). The main effect of increasing the concentration of agonist was to increase the open probability within bursts, with saturating concentrations of full agonists, such as acetylcholine, leading to bursts where the open probability exceeded 0.90 (i.e. the channel was open for more than 90% of the duration of the burst). Spontaneous openings of β_Anc_ homopentamers also occur in bursts, and despite the absence of agonist, activation appears efficient, with the probability of being open within a burst also exceeding 0.90. The kinetic structure of bursts of spontaneous β_Anc_ openings also resembles that of the agonist-activated AChR, with bursts containing several types of closings, the briefest of which are reminiscent of classic “*nachschlag* shuttings”. This is significant because it was originally proposed that the duration of *nachschlag* shuttings was related to agonist efficacy (22). However, subsequent work showed that *nachschlag* shuttings were independent of agonist, which necessitated the introduction of an additional closed state appended to the agonist-activated open state in early schemes of AChR activation (41). This latter finding was some of the impetus for refined schemes, where the additional closed states preceded channel opening and were referred to as “flipped” or “primed” (23, 24). Our kinetic analysis has shown that the spontaneous activity of β_Anc_ fits an analogous scheme, containing an intermediate closed state that also precedes channel opening, but where the agonist binding steps have been omitted due to the absence of agonist. Evidently, these functional hallmarks of AChR activation do not arise from the complex heteropentameric architecture of the muscle-type receptor, nor do they depend upon the presence of agonist. Instead, they are fundamental properties preserved and encoded in the reconstructed amino acid sequence of β_Anc_.

Based upon the rates of open-channel block by acetylcholine and the well-known open-channel blocker, QX-222, the structure of the open pore in β_Anc_ homopentamers and wild-type AChRs also appears to be similar. This is not surprising, since open-channel block is determined primarily by interactions between the blocking molecule and the second transmembrane segments (i.e. M2) of each AChR subunit (26–28, 42, 43). The M2 segment, which lines the channel pore at its narrowest constriction, is a well-known determinant of single-channel conductance and block, and is one of the more conserved regions across all pentameric ligand-gated ion channel subunits (2). Accordingly, this site is largely conserved in β_Anc_ (9), thus the chemical composition and profile of the open pore in β_Anc_ homopentamers is expected to resemble that of the open pore in the wild-type AChR.

A degree of uncertainty is inherent in the reconstruction of any ancestral protein, and to solidify evolutionary conclusions it is important to assess whether the functions of putative ancestors are robust to these uncertainties. We have shown that an alternate ancestral β-subunit (β_AncS_), with 67 substitutions and 6 indels relative to β_Anc_, was still able to form homomers that opened spontaneously. Furthermore, spontaneous openings of β_AncS_ homomers also occurred in bursts that contained brief closings (i.e., “*nachschlag* shuttings”), and had an open probability that exceeded 0.90. These findings demonstrate that the ability of β_Anc_ to form spontaneously opening homomers is robust to substantial variations in the inferred ancestral β-subunit amino acid sequence. Thus, the ability of these reconstructed ancestral subunits to form spontaneously opening homomers is deeply embedded in their structure and evolutionary history.

The kinetic behaviour of β_AncS_ homomers was more complex than originally observed with β_Anc_. This increased complexity necessitated the expansion of the original scheme used to fit β_Anc_ to include two additional priming steps. Both the original β_Anc_ and expanded β_AncS_ schemes parallel the scheme used to describe the muscle-type AChR, which included two priming steps (i.e. “singly” and “doubly” primed) that each correlated with conformational changes around the two agonist-binding sites (24). In the case of β_AncS_, which lacks agonist-binding sites, the simplest interpretation is that the different levels of priming correlate with conformational changes occurring within individual subunits. Although they are labelled as “singly”, “doubly”, and “triply” primed, assuming that β_AncS_ also forms homopentamers, the three priming steps in β_AncS_ homomers could represent the three most terminal priming steps, where three, four, or five β_AncS_ subunits are primed. Within this framework, openings from β_AncS_ channels with zero, one, or two subunits primed, are presumably unstable, and thus not observed in our single-channel recordings. In an alternate scenario, “singly” and “doubly” primed could refer to β_AncS_ channels with one or two primed subunits, respectively. While “triply” primed could refer to channels with three or more primed subunits, but where openings from β_AncS_ channels with three, four, or five primed subunits are indistinguishable. Applied to β_Anc_, these interpretations suggest that either (1) openings from only the terminal priming step, where all five β_Anc_ subunits are primed, are stable enough to be observed in β_Anc_ homopentamers, or (2) openings can occur with fewer than five primed subunits, but these openings are kinetically indistinguishable. Regardless of the interpretation, priming is an important step in the activation of both β_AncS_ and β_Anc_ homopentamers.

Modern mechanisms of AChR activation include intermediate closed states, which place the roots of agonism at a stage in the activation process that precedes channel opening (13, 23, 24). A consequence of these mechanisms is that the ultimate opening and closing rates of the channel are independent of agonist. Here, we have shown that additional single-channel hallmarks of AChR function are also independent of agonist, as they occur in spontaneously opening homopentameric channels that are devoid of agonist-binding sites, and which are formed from reconstructed ancestral AChR β-subunits. Often overlooked, the β-subunit is the least conserved of the four AChR subunits, and is the only subunit that does not contribute residues to the two AChR agonist-binding sites. Despite these considerations, hallmarks of AChR function remain deeply embedded in β-subunit sequence, structure, and evolutionary history. Given that these functional hallmarks are independent of agonist, it is tempting to speculate that they predate agonism, and thus that agonism evolved subsequently as an additional layer of regulation in this family of pentameric ion channels.

## Acknowledgements

C.J.G.T. was funded in part by an Ontario Graduate Scholarship, while J.R.E. was funded in part by a Natural Sciences and Engineering Research Council of Canada (NSERC) CREATE Scholarship. C.J.B.d.C. acknowledges grants from NSERC (RGPIN-2016-04801), the Canada Foundation for Innovation (34475), and the Canadian Institutes of Health Research (377068).

## Author Contributions

C.J.G.T. acquired and analyzed all electrophysiological data, except the data for the electrical fingerprinting, which was recorded and analyzed by R.M.S., C.J.G.T, and C.J.B.d.C. C.J.G.T., F.J.P.A. and J.R.E. acquired some of the initial recordings containing β_Anc_ homopentamers. G.D.M. reconstructed β_AncS_ and acquired β_AncS_ electrophysiological data. C.J.G.T. and C.J.B.d.C. interpreted the data and wrote the manuscript. C.J.B.d.C. supervised the project.

## Conflict of Interest

The authors declare that they have no conflicts of interest with the contents of this article.

## SUPPLEMENTARY INFORMATION

**Figure S1:**
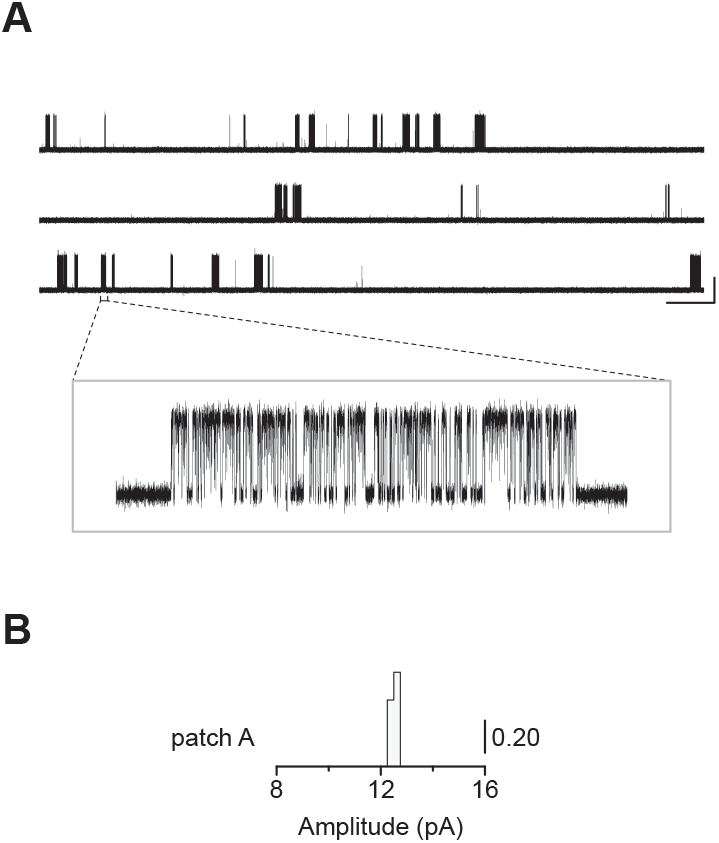
Single-channel recordings of the human adult muscle-type acetylcholine receptor exhibit homogeneous burst behaviour. (A) Representative continuous recording from a cell-attached patch where cells were transfected with cDNAs encoding human muscle-type α-, β-, δ-, and ε-subunits at a cDNA ratio of 2:1:1:1 (α:β:δ:ε). Activity was recorded in the presence of 30 μM acetylcholine, with an applied voltage of –120 mV. Continuous trace (top) was digitally filtered to 5 kHz (scale bar = 2 s, 10 pA), while inset burst (boxed; bottom) was filtered to 10 kHz where box itself represents 300 ms and 25 pA. Openings are shown as upward deflections. (B) Event-based amplitude histogram for single-channel bursts for the patch shown in panel “A”. The height of the bins was normalized to the total number of bursts in the patch (29 total), with the scale bar representing 0.20 of the total number of bursts.

**Figure S2:**
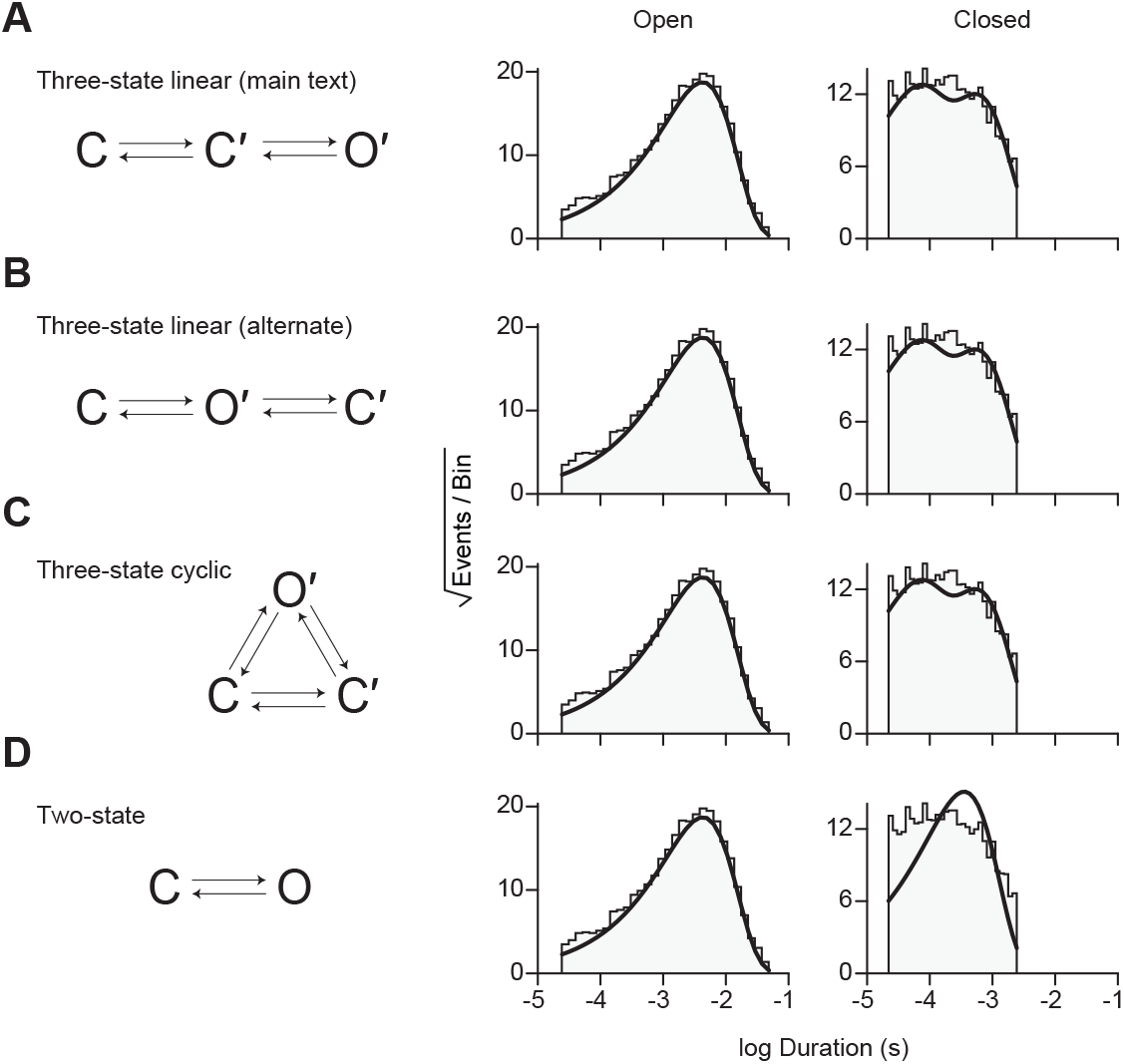
Kinetic fitting of alternate schemes describing spontaneous single-channel activity of β_Anc_ homomers. Each scheme is shown on the left with the same open and closed duration histograms shown on the right. In each case the resulting fit of the same data using the various schemes is overlaid (solid black lines). (A) The three-state linear scheme described in the main text is reproduced here for comparison with (B) an alternate three-state linear scheme, as well as (C) the three-state cyclic scheme. (D) A two-state scheme with a single closed state and a single open state was also fit to validate the need for a second closed state.

**Figure S3:**
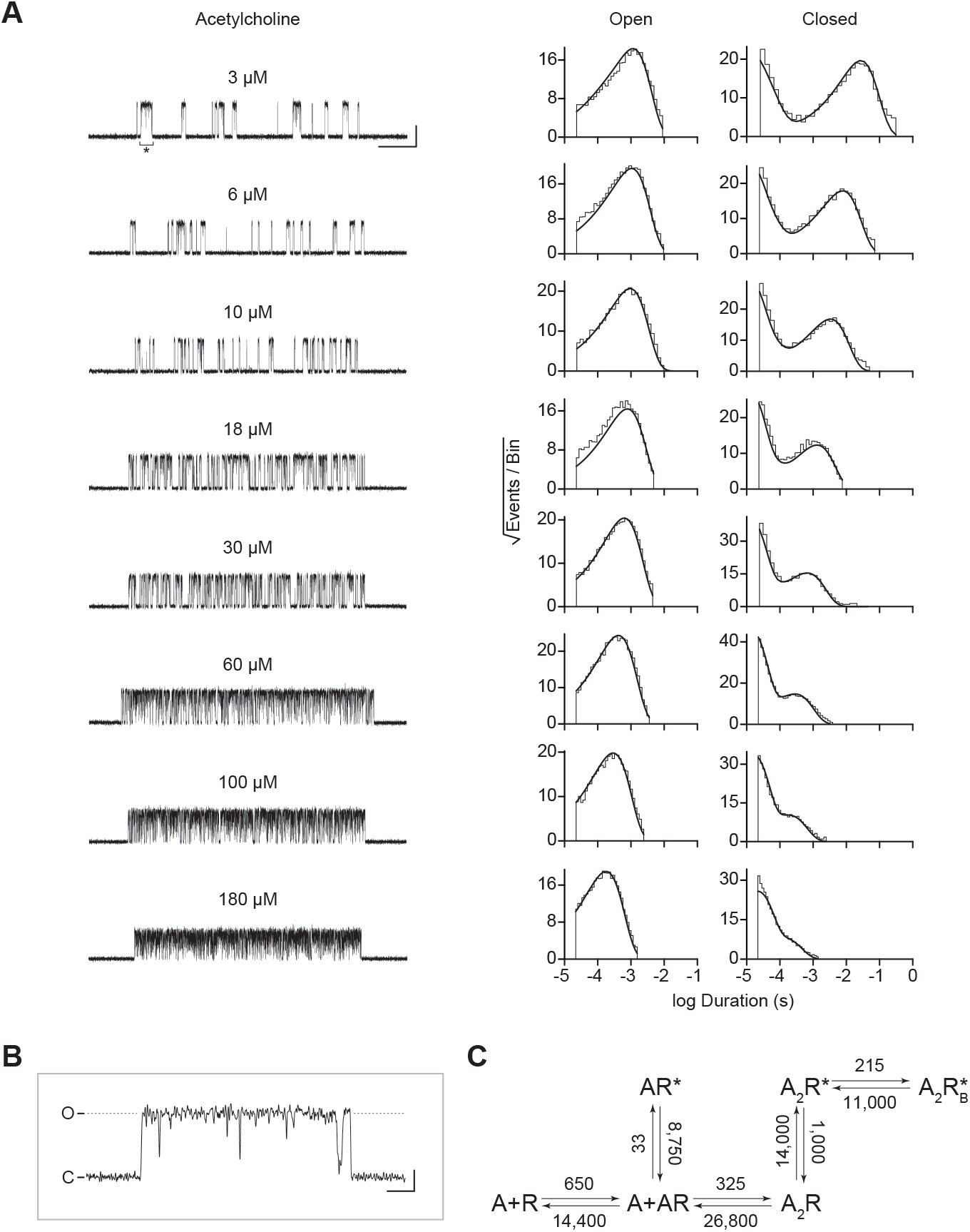
Single-channel kinetics of the human adult muscle-type acetylcholine receptor. (A) Representative bursts from single-channel recordings of the wild-type AChR over a full acetylcholine concentration range (left). Recordings were acquired in the cell-attached patch configuration, with an applied voltage of –120 mV. Openings are upward deflections, bursts are filtered to 10 kHz, and the scale bar represents 25 ms and 10 pA. Corresponding open and closed duration histograms are presented for each concentration (right), with overlaid fits (solid line) from global fitting of the entire data set. (B) Representative openings interrupted by “*nachschlag* shuttings” in the presence of 3 μM acetylcholine (see asterisk in “A”), where “C” represents the closed, baseline current, and “O” represents the current through a single open conducting AChR. Scale bar in “B” represents 1 ms and 5 pA. (C) Modified del Castillo and Katz kinetic scheme (see reference no. 19 of main text) for the muscle-type AChR, containing two agonist binding steps, and a single blocked state flanking the doubly-liganded open state (Scheme 4). Estimates of the rate constants from global fitting of the entire data set are shown, with error estimates presented in Table S2. Global kinetic fits were performed on three individual recordings for each concentration of acetylcholine, from at least two separate transfections, corresponding to 24 total patches for the entire global fit.

**Figure S4:**
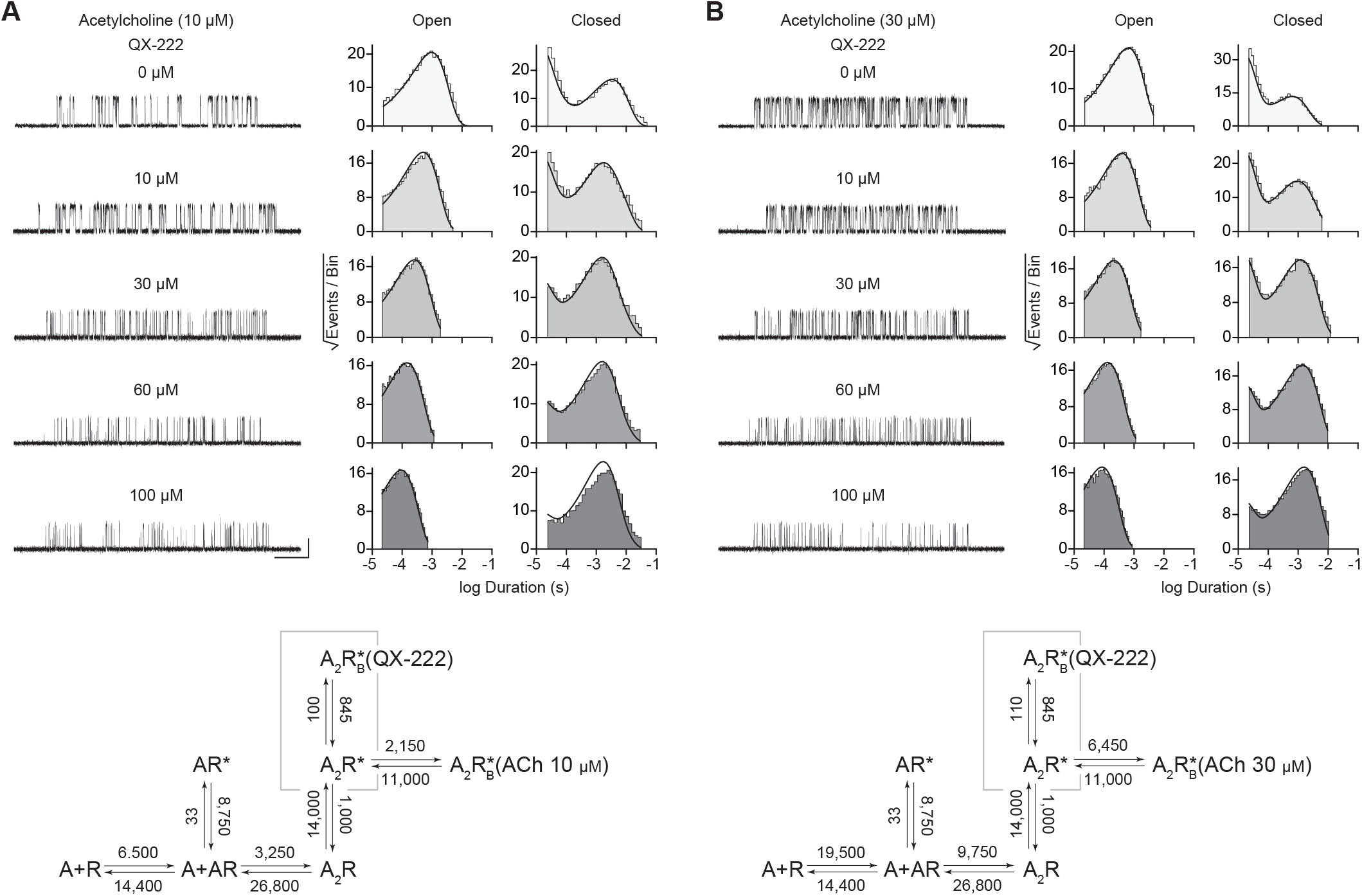
Kinetics of QX-222 block of the human adult muscle-type acetylcholine receptor. Single-channel recordings were obtained with increasing concentrations of QX-222, while the concentration of agonist (acetylcholine; ACh) was kept constant at either (A) 10 μM or (B) 30 μM. Single channel recordings were obtained in the cell-attached configuration, with an applied voltage of –120 mV, where openings are upward deflections, and traces filtered to 10 kHz. Corresponding open and closed duration histograms are presented for all QX-222 concentrations with overlaid fits (solid line) from global fitting of the entire data set. Kinetic scheme is the same presented in Figure S3, but with an additional open/blocked state (corresponding to QX-222 block) flanking the doubly-liganded open state (Scheme 5). All rates, except those describing QX-222 block, were fixed to that at the indicated acetylcholine concentration as estimated from original QX-222-free fits in Figure S3. For both concentrations of acetylcholine, estimates of the rate constants from global fitting of each data set are shown, with error estimates presented in Table S2. Global kinetic fits were performed on three individual recordings for each concentration of QX-222 at both concentrations of ACh, and from at least two separate transfections, corresponding to 15 total patches for each global fit.

**Figure S5:**
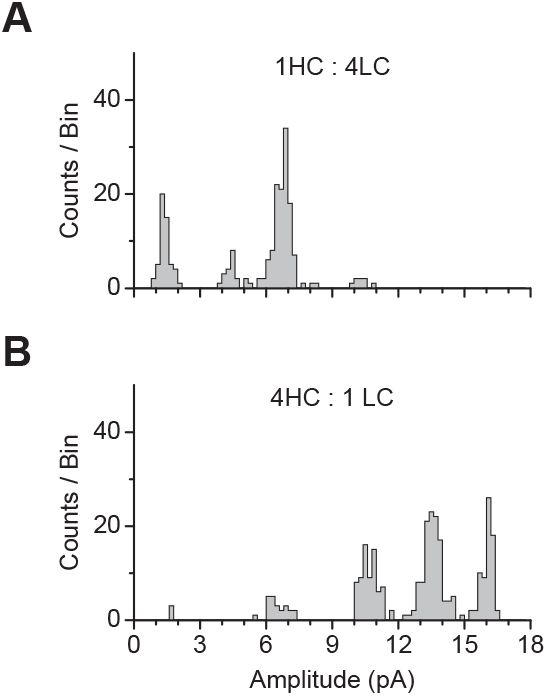
Event-based amplitude histograms derived from a representative single patch where cells were cotransfected with cDNAs encoding for high-conductance (HC) and low-conductance (LC) variants of β_Anc_ at a ratio (mass:mass) of (A) 1HC:4LC or (B) 4HC:1LC. In each case, cotransfection of HC and LC β_Anc_ subunits led to a distribution of amplitudes that segregated into distinct amplitude classes, where the proportion of events in each class was biased by the relative proportion of each type of cDNA transfected. These two patches with overlapping amplitude classes were combined to produce the plot in Figure 5D.

**Table S1:** Single-channel kinetics of spontaneously opening β_Anc_ homomers.

| Homomer | $k_{+1}'$ | $k_{-1}'$ | $K'$ | $\beta_1$ | $\alpha_1$ | $\Theta_1$ | $k_{+B}$ | $k_{-B}$ | $K_B$<br>($\mu\text{M}$) |
| --- | --- | --- | --- | --- | --- | --- | --- | --- | --- |
| No Agonist | 3,900 | 7,600 | 1.95 | 10,000 | 310 | 32.26 | N/A | N/A | N/A |
| (3 patches) | (110) | (450) |  | (300) | (3) |  |  |  |  |
| Acetylcholine | 3,500 | 5,400 | 1.54 | 8,000 | 330 | 24.24 | 170* | 62,000 | 364.71 |
| (15 patches) | (85) | (265) |  | (200) | (3) |  | (1.8) | (450) |  |
| QX-222 | 4,000 | 8,000 | 2.00 | 10,500 | 320 | 32.81 | 95* | 1,400 | 14.74 |
| (15 patches) | (110) | (440) |  | (300) | (3) |  | (0.5) | (7) |  |
| Acetylcholine<br>(Constrained) | 3,900 | 7,600 | 1.95 | 10,000 | 310 | 32.26 | 170* | 60,800 | 357.65 |
| (15 patches) |  |  |  |  |  |  | (1.5) | (410) |  |
| QX-222<br>(Constrained) | 3,900 | 7,600 | 1.95 | 10,000 | 310 | 32.26 | 95* | 1,400 | 14.74 |
| (15 patches) |  |  |  |  |  |  | (0.5) | (6.5) |  |

*Note:* Rate constants were estimated from fitting Scheme 1 or Scheme 2 (below) presented in Figures 3 and 4, respectively. Data were globally fit (number of patches indicated in each case) over a range of acetylcholine/QX-222 concentrations, with rate constants and associated errors (parentheses) estimated by MIL (see Experimental Procedures). Priming ( $K'$ ), gating ( $\theta_1$ ) and blocking ( $K_B$ ) equilibrium constants represent $k_{-1}'/k_{+1}'$ , $\beta_1/\alpha_1$ , and $k_{-B}/k_{+B}$ , respectively. Association rate constants (\*) ( $k_{+B}$ ) are presented in units of $\mu\text{M}^{-1}\cdot\text{s}^{-1}$ , while remaining rate constants are presented in units of $\text{s}^{-1}$ . Constrained rates (presented in grey) were held constant at the values determined from the agonist free dataset, while the blocking rate constants were estimated.

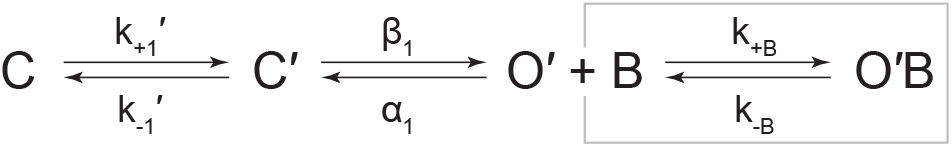
Scheme 1 & Scheme 2 (includes boxed)

**Table S2:** Kinetics of acetylcholine activation and QX-222 block of human adult muscle-type acetylcholine receptors.

| WT | $k_{+1}$ | $k_{-1}$ | $K_1$<br>($\mu\text{M}$) | $k_{+2}$ | $k_{-2}$ | $K_2$<br>($\mu\text{M}$) | $\beta_1$ | $\alpha_1$ | $\Theta_1$ | $\beta_2$ | $\alpha_2$ | $\Theta_2$ | $k_{+B}$ (ACh) | $k_{-B}$ (ACh) | $K_{B(\text{ACh})}$<br>($\mu\text{M}$) | $k_{+B}$<br>(QX-222) | $k_{-B}$<br>(QX-222) | $K_B$<br>(QX-222)<br>($\mu\text{M}$) |
| --- | --- | --- | --- | --- | --- | --- | --- | --- | --- | --- | --- | --- | --- | --- | --- | --- | --- | --- |
| ACh<br>(24 pat.) | 650*<br>(35) | 14,400<br>(1000) | 22.12 | 325*<br>(20) | 26,500<br>(400) | 81.54 | 33<br>(2.5) | 8,750<br>(650) | 3.77<br>E-03 | 14,000<br>(450) | 1,000<br>(10) | 14 | 215*<br>(3) | 110,000<br>(800) | 511.63 | N/A | N/A | N/A |
| QX-222<br>(10 $\mu\text{M}$ ACh)<br>(15 pat.) | 6,500 | 14,400 | 2.21 | 3,250 | 26,500 | 8.15 | 33 | 8,750 | 3.77<br>E-03 | 14,000 | 1,000 | 14 | 2,150 | 110,000 | 51.16 | 100*<br>(0.5) | 845<br>(5.5) | 8.45 |
| QX-222<br>(30 $\mu\text{M}$ ACh)<br>(15 pat.) | 19,500 | 14,400 | 0.74 | 9,750 | 26,500 | 2.72 | 33 | 8,750 | 3.77<br>E-03 | 14,000 | 1,000 | 14 | 6,450 | 110,000 | 17.05 | 110*<br>(0.5) | 845<br>(4.5) | 7.68 |

*Note:* Rate constants were estimated from fitting Scheme 4 or Scheme 5 (below) presented in Figure S3 or Figure S4, respectively. Where “A” represents agonist, and “R”, “R\*”, and “R\*B” represent the closed, open, and open-blocked states of the human adult muscle-type acetylcholine receptor. Data were globally fit (number of patches indicated in each case) over a range of QX-222 concentrations with fixed concentration of acetylcholine (ACh: 10 $\mu\text{M}$ or 30 $\mu\text{M}$ ), with rate constants and associated errors (parentheses) estimated with MIL (see Experimental Procedures). Apparent binding ( $K_n$ ), apparent gating ( $\theta_n$ ) and blocking ( $K_B$ ) equilibrium constants represent $k_{-n}/k_{+n}$ , $\beta_n/\alpha_n$ , and $k_{-B}/k_{+B}$ , respectively. Association rate constants (\*; $k_{+1}$ , $k_{+2}$ , $k_{+B}$ ) are presented in units of $\mu\text{M}^{-1}\cdot\text{s}^{-1}$ , while remaining rate constants are presented in units of $\text{s}^{-1}$ . Constrained rates (grey) were held constant to the rates estimated from fitting of wild-type (WT) in the absence of QX-222, allowing the QX-222 blocking rates to be estimated independently.

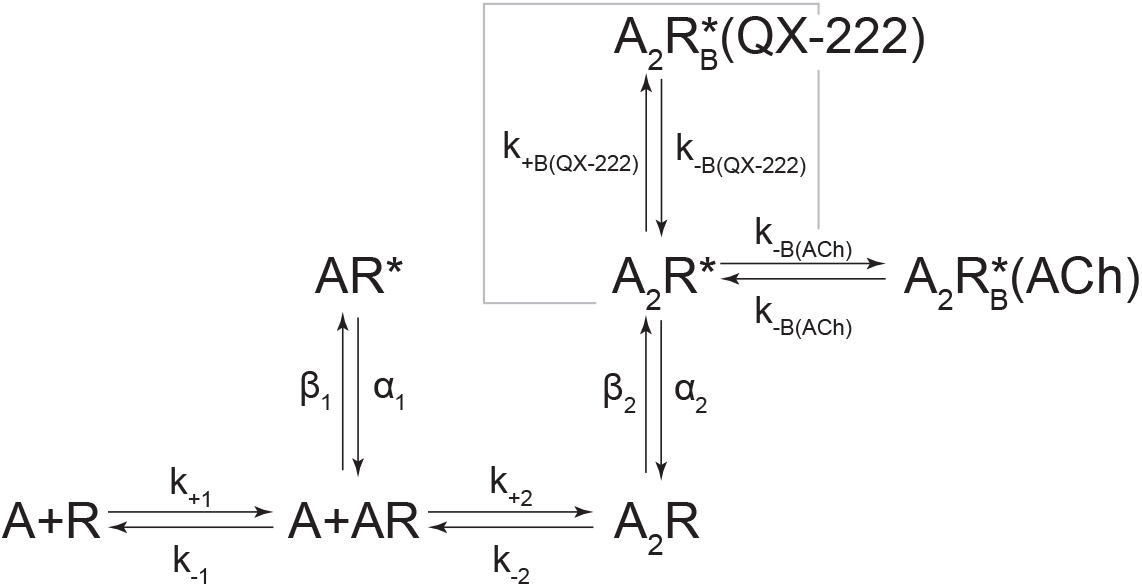
Scheme 4 & Scheme 5 (includes boxed)

**Table S3:**
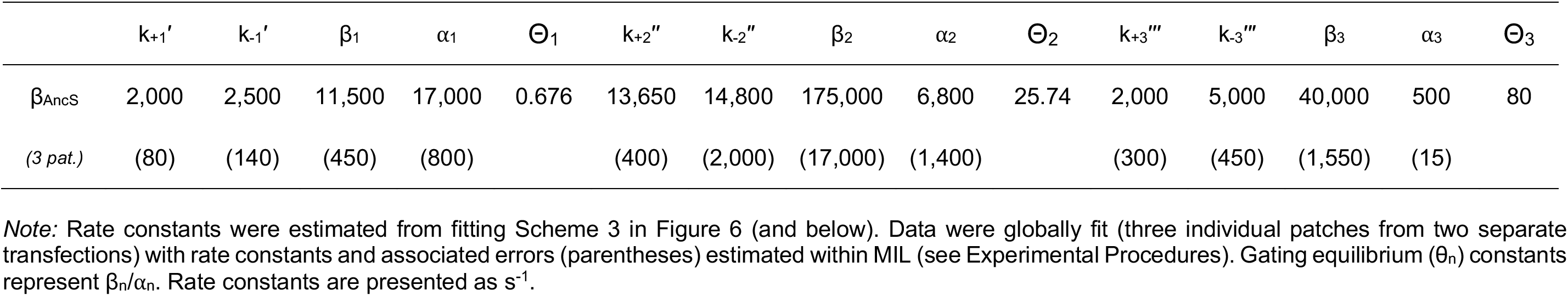
Single-channel kinetics of spontaneously opening β_AncS_ homomers.

| | $k_{+1}'$ | $k_{-1}'$ | $\beta_1$ | $\alpha_1$ | $\Theta_1$ | $k_{+2}''$ | $k_{-2}''$ | $\beta_2$ | $\alpha_2$ | $\Theta_2$ | $k_{+3}'''$ | $k_{-3}'''$ | $\beta_3$ | $\alpha_3$ | $\Theta_3$ |
| --- | --- | --- | --- | --- | --- | --- | --- | --- | --- | --- | --- | --- | --- | --- | --- |
| $\beta_{\text{AncS}}$ | 2,000 | 2,500 | 11,500 | 17,000 | 0.676 | 13,650 | 14,800 | 175,000 | 6,800 | 25.74 | 2,000 | 5,000 | 40,000 | 500 | 80 |
| (3 pat.) | (80) | (140) | (450) | (800) |  | (400) | (2,000) | (17,000) | (1,400) |  | (300) | (450) | (1,550) | (15) |  |

*Note:* Rate constants were estimated from fitting Scheme 3 in Figure 6 (and below). Data were globally fit (three individual patches from two separate transfections) with rate constants and associated errors (parentheses) estimated within MIL (see Experimental Procedures). Gating equilibrium ( $\theta_n$ ) constants represent $\beta_n/\alpha_n$ . Rate constants are presented as $\text{s}^{-1}$ .

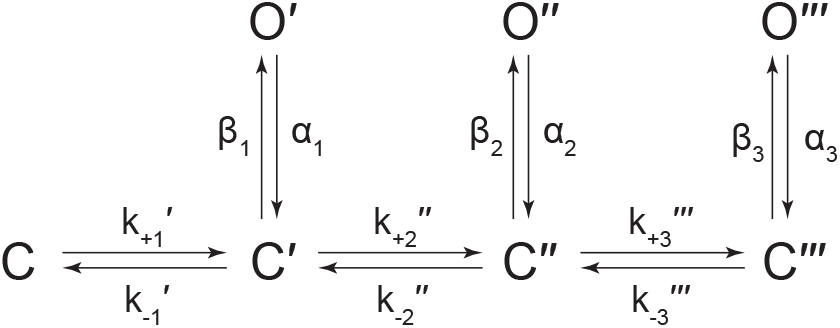
Scheme 3.

### Description of Source Data files

**Source Data - Figure 3.zip**

- Source data for Figure 3. Detected single-channel event durations of spontaneously opening β_Anc_ homomers. Compressed file includes three TAC event files (*.evt format) of the single-channel detections for the three recordings used in the presented kinetic analysis, as well as the associated R scripts (*.txt format) for defining and sorting bursts. *(6 files total)*

**Source Data - Figure 4.zip**

- Source data for Figure 4. Detected single-channel event durations of spontaneously opening β_Anc_ homomers in the absence (AgonistFree) and presence of increasing concentrations of acetylcholine (ACh; 10 μM, 30 μM, 60 μM and 100 μM) or QX-222 (QX; 10 μM, 30 μM, 60 μM and 100 μM). Compressed file includes 27 TAC event files (*.evt format) of the single-channel detections for the three recordings for each condition (fileA, fileB, fileC in each case) in the presented kinetic analysis, as well as the associated R scripts (*.txt format) for defining and sorting bursts.

(54 files total)

**Source Data - Figure 6.zip**

- Source data for Figure 6. Detected single-channel event durations of spontaneously opening β_AncS_ homomers. Compressed file includes three TAC event files (*.evt format) of the single-channel detections for the three recordings used in the presented kinetic analysis, as well as the associated R scripts (*.txt format) for defining and sorting bursts. *(6 files total)*

**Source Data - Figure S3.zip**

- Source data for Figure S3. Detected single-channel event durations for the wild-type adult muscle acetylcholine receptor in the presence of increasing concentrations of acetylcholine (ACh; 3 μM, 6μM, 10 μM, 18 μM, 30 μM, 60 μM, 100 μM and 180 μM). Compressed file includes 24 TAC event files (*.evt format) of the single-channel detections for the three recordings for each acetylcholine concentration (fileA, fileB, fileC in each case) in the presented kinetic analysis, as well as the associated R scripts (*.txt format) for defining and sorting bursts.

(48 files total)

**Source Data - Figure S4.zip**

- Source data for Figure S3. Detected single-channel event durations for QX-222 block of the wild-type adult muscle acetylcholine receptor. Recordings were in the presence of either 10 μM or 30 μM acetylcholine (ACh), and increasing concentrations of QX-222 (QX222; 10 μM, 30 μM, 60 μM and 100 μM). Compressed file includes 30 TAC event files (*.evt format) of the single-channel detections for three recordings for each condition (fileA, fileB, fileC in each case) included in the kinetic analysis, as well as the associated R scripts (*.txt format) for defining and sorting bursts.

(60 files total)

## Notes

### Competing Interest Statement

The authors have declared no competing interest.

